# Structure, Asymmetry and Segmentation of the Human Parietal Aslant and Vertical Occipital Fasciculi

**DOI:** 10.1101/252825

**Authors:** Sandip S Panesar, Joao Tiago A Belo, Fang-Cheng Yeh, Juan C Fernandez-Miranda

## Abstract

We previously proposed a bipartite ‘dorsal-ventral’ model of human arcuate fasciculus (AF) morphology. This model does not, however, account for the ‘vertical,’ temporoparietal subdivision of the AF described in earlier dissection and tractographic studies. In an effort to address the absence of the vertical AF (VAF) within the ‘dorsal-ventral’ model, we conducted a dedicated tractographic and white-matter dissection study of this tract and another short, vertical, posterior-hemispheric fascicle: the vertical occipital fasciculus (VOF). We conducted atlas-based, non-tensor, deterministic tractography in 30 single subjects from the Human Connectome Project database and verified our results using an average diffusion atlas comprising 842 separate normal subjects. We also performed white-matter dissection in 4 cadaveric hemispheres. Our tractographic results demonstrate that the VAF is in fact a bipartite system connecting the ventral-parietal and ventral-temporal regions, with variable connective and no volumetric lateralization. The VOF is a nonlateralized, non-segmented system connecting lateral occipital areas with basal-temporal regions. Importantly, the VOF was distinctly dissociated from the VAF. As the VAF demonstrates no overall connective or volumetric lateralization, we postulate its distinction from the AF system and propose its renaming to the ‘parietal aslant tract,’ (PAT) with unique dorsal and ventral subdivisions. Our tractography results were supported by diffusion atlas and white matter dissection findings.

## 1 Introduction

The anatomy, classification, and even existence of posterior hemispheric association white matter tracts have been subject to controversy in the anatomical literature. Early tractographic and fiber dissection descriptions of the arcuate fasciculus (AF) included a subcomponent known as the ‘vertical arcuate fasciculus’, which comprised one of three AF subcomponents (Catani, Howard, Pajevic, & Jones, 2002; Catani, Jones, & ffytche, 2005; Fernandez-Miranda et al., 2008). The ‘vertical AF’ (VAF) was described as a short, vertically-oriented tract originating within the posterior temporal lobe and terminating within the inferior parietal cortex. Along with the fronto-parietal horizontal subcomponent, it formed the ‘superficial-AF’ which lay superficial to the temporo-frontal perisylvian component to comprise the ‘AF’ in its entirety. Proponents of this original description (Catani & Thiebaut de Schotten, 2008; Martino, da Silva-Freitas, et al., 2013; Martino, De Witt Hamer, et al., 2013; Martino & Garcia-Porrero, 2013) regard the AF as part of the greater superior longitudinal fasciculus (SLF) system, with the AF referred to as the ‘perisylvian-SLF’ or ‘SLF-IV.’ More recent fiber dissection and tractographic studies have questioned this description, proposing instead the AF as a purely temporo-frontal tract, with a ‘dorsal-ventral’ arrangement and without any temporo-parietal or fronto-parietal connectivity (Fernandez-Miranda et al., 2015; Glasser & Rilling, 2008; Lawes et al., 2008; Rilling et al., 2008). This view contradicts postulations of the AF and SLF belonging to the same fiber system, as in this model the AF, but not the SLF, is strongly leftward lateralized in connectivity and volumetry. Tractographic studies have demonstrated the SLFs rightward-dominant connectivity and volumetric lateralization (Thiebaut de Schotten et al., 2011; Wang et al., 2016). The dorsal-ventral AF argument leaves the previously described VAF unaccounted for. The relatively recent tractography-derived description of the frontal-aslant tract (FAT) (Catani et al., 2013) demonstrates that short, low-volume white matter systems that were previously obscured due to limitations of white fiber dissection techniques can be reliably studied by tractography. Furthermore, the limitations of early tractographic techniques (i.e. DTI), including inability to trace crossing fiber pathways (Farquharson et al., 2013), have been successfully addressed by newer methods including the non-tensor diffusion spectrum imaging (DSI) algorithm (Yeh, Wedeen, & Tseng, 2010). As such, tractography provides an excellent medium to study shorter white matter populations including the previously described VAF.

Another described tract known as the vertical occipital fasciculus (VOF) is a connection within the extreme posterior hemisphere that was first described towards the end of the 19^th^ century. Its description disappeared from neuroanatomical literature until the introduction of autoradiographic and tractographic techniques from the 1970s onwards (Weiner, Yeatman, & Wandell, 2016; Yeatman et al., 2014). A dedicated study by Yeatman et al. (2013) defined the anatomy of the posterior visual word-form area, and tractographically reproduced this bundle using tensor methods. The authors described a tract ascending from the occipitotemporal sulcus lateral to the inferior longitudinal fasciculus (ILF) and terminating within the parieto-occipital association cortices. Nevertheless, relative to the well-known large association fascicles, little data exists pertaining to the VOFs exact anatomy.

With these considerations we conducted a dedicated tractographic and white matter dissection study into the morphology, volumetry, and connectivity of the VAF and VOF in an effort to address several issues: 1.) The missing VAF in the contemporary ‘dorsal-ventral’ AF model. 2.) Apply a deterministic non-tensor tractographic algorithm to address the issue of discreet or unified VAF and VOF. 3.) Specifically analyze the lateralization of the VAF to determine whether it yielded further evidence to the argument regarding dissociation of the AF and SLF systems in the contemporary lexicon.

## 2. Methods

Three principal methods were used in this study, firstly a dedicated subject-specific tractography study was conducted in 30 individual subjects. Then, we utilized a diffusion tractography atlas consisting of averaged diffusion data from 842 individual subjects for verification of results. Finally, both tractographic methods were complemented by a white matter dissection study in cadaveric brains.

### 2.1 Participants

We conducted a subject-specific deterministic fiber tractography study in 30 right-handed, neurologically-healthy male and female subjects, age range 23-35. The data were from the Human Connectome Project (HCP) online database (WU-Minn Consortium (Principal Investigators: David van Essen and Kamil Ugurbil; 1U54MH091657) funded by the 16 NIH institutes and centers that support the NIH Blueprint for Neuroscience Research and by the McDonnell Center for Systems Neuroscience at Washington University. Likewise, data from 842 individual HCP subjects were utilized to compile the averaged diffusion atlas.

### 2.2 Image Acquisition and Reconstruction

The HCP diffusion data for individual subjects were acquired using a Siemens 3T Skyra system, with a 32-channel head coil (Siemans Medical, Erlangen, Germany). A multishell diffusion scheme was used, and the *b* values were 1000, 2000, and 3000 s/mm^2^. The number of diffusion sampling directions were 90, 90 and 90, respectively. The in-plane resolution and slice thickness were both 1.25mm (TR = 5500 ms, TE = 89 ms, resolution = 1.25mm x 1.25mm, FoV = 210mm x 180mm, matrix = 144 x 168). The DSI data were reconstructed using the generalized q-sampling imaging approach (Yeh et al., 2010)using a diffusion distance ratio of 1.2 as recommended by the original study.

### 2.3 HCP 842 Atlas

A total of 842 participants from the HCP database were used to construct the atlas. The image acquisition parameters are identical to those in 2.2. The diffusion data were reconstructed and warped to the Montreal Neurological Institute (MNI) space using *q-*space diffeomorphic reconstruction (Yeh & Tseng, 2011) with a diffusion sampling length ratio of 1.25 and the output resolution was 1 mm. The group average atlas was then constructed by averaging the reconstructed data of the 842 individual subjects within the MNI space.

### 2.4 Fiber Tracking and Analysis

We performed deterministic fiber tracking using DSI Studio software (http://dsi-studio.labsolver.org), which utilizes a generalized streamline fiber tracking method (Yeh, Verstynen, Wang, Fernandez-Miranda, & Tseng, 2013). Parameters selected for fiber tracking included a step size of 0.2 mm, a minimum fiber length of 20mm and a turning angle threshold of 60°. For progression locations containing >1 fiber orientation, fiber orientation most congruent with the incoming direction and turning angle <60° was selected to determine subsequent moving direction. Each progressive voxels moving directional estimate was weighted by 20% of the previous voxels incoming direction and by 80% if its nearest fiber orientation. This sequence was repeated to create fiber tracts. Termination of the tracking algorithm occurred when the quantitative anisotropy (QA) (Yeh et al., 2013) dropped below a subject-specific, pre-selected threshold value of between 0.02-0.08, when fiber tract continuity no longer met the progression criteria, or when 100,000 tracts were generated. We pre-selected QA termination threshold by analyzing the number of false continuations generated within each subjects’ dataset and chose the compromise value that allowed optimal anatomical detail with minimal noise. Likewise, we selected a smoothing parameter of 50% for the same reason stated previously.

To generate the desired tracts, we employed an atlas-based approach (Fernandez-Miranda et al., 2015; Panesar et al., 2017; Wang et al., 2016), along with manually placed regions of interest (ROI) and regions of avoidance (ROA). We utilized the Automated Anatomical Labelling atlas (AAL) (Tzourio-Mazoyer et al., 2002) to select cortical seeding regions. Briefly, the AAL regions were warped to each individual subjects’ diffusion matrix using the linear and non-linear registration algorithm feature in DSI Studio. This feature allows optimal alignment of cortical atlas regions to each subjects’ diffusion map, requiring little-to-no manual manipulation of regions. For the VAF we chose all parietal regions, as parcellated by the AAL: precuneus (Pc), superior parietal lobule (SPL), and inferior parietal lobules (dorsal and ventral aspects of the supramarginal gyrus (dSmG; vSMG) and angular gyrus (AG)). A rectangular axial ROI was placed ventral to the seeding regions to select only fibers passing dorsal-ventral (i.e. parietal-temporal/parietal-occipital), and an additional cross-shaped rectangular ROA was utilized, with one portion lying in the coronal plane and one in the sagittal plane. The coronal ROA was to prevent any anteriorly travelling fibers from being generated, and the sagittal ROA was used to prevent any of the commissural fibers from being generated. VOF fibers were generated separately. Three superior occipital regions gyri were chosen as per the AAL atlas (superior occipital gyrus (SO); middle occipital gyrus (MO); cuneus (Cu)). The same rectangular ROI was utilized, but positioned ventral relative to its positioning for the vertical-AF, at the position of the ventral temporo-occipital regions (Figure 1A, B). The process was repeated for both right and left hemispheric vertical-AFs and VOFs. The chosen atlas-based approach to fiber tracking was to ensure consistency of methodology across subjects, with minimal permitted *a priori* or inter-user variability. Once 100,000 fiber tracts had been generated, we manually selected fibers travelling parieto-temporally, parieto-occipitally, occipito-occipitally or occipito-temporally for further analysis. We subsequently analyzed the dorsal and ventral morphology and connectivity of each bundle for connectivity analysis.

**1A-D.**
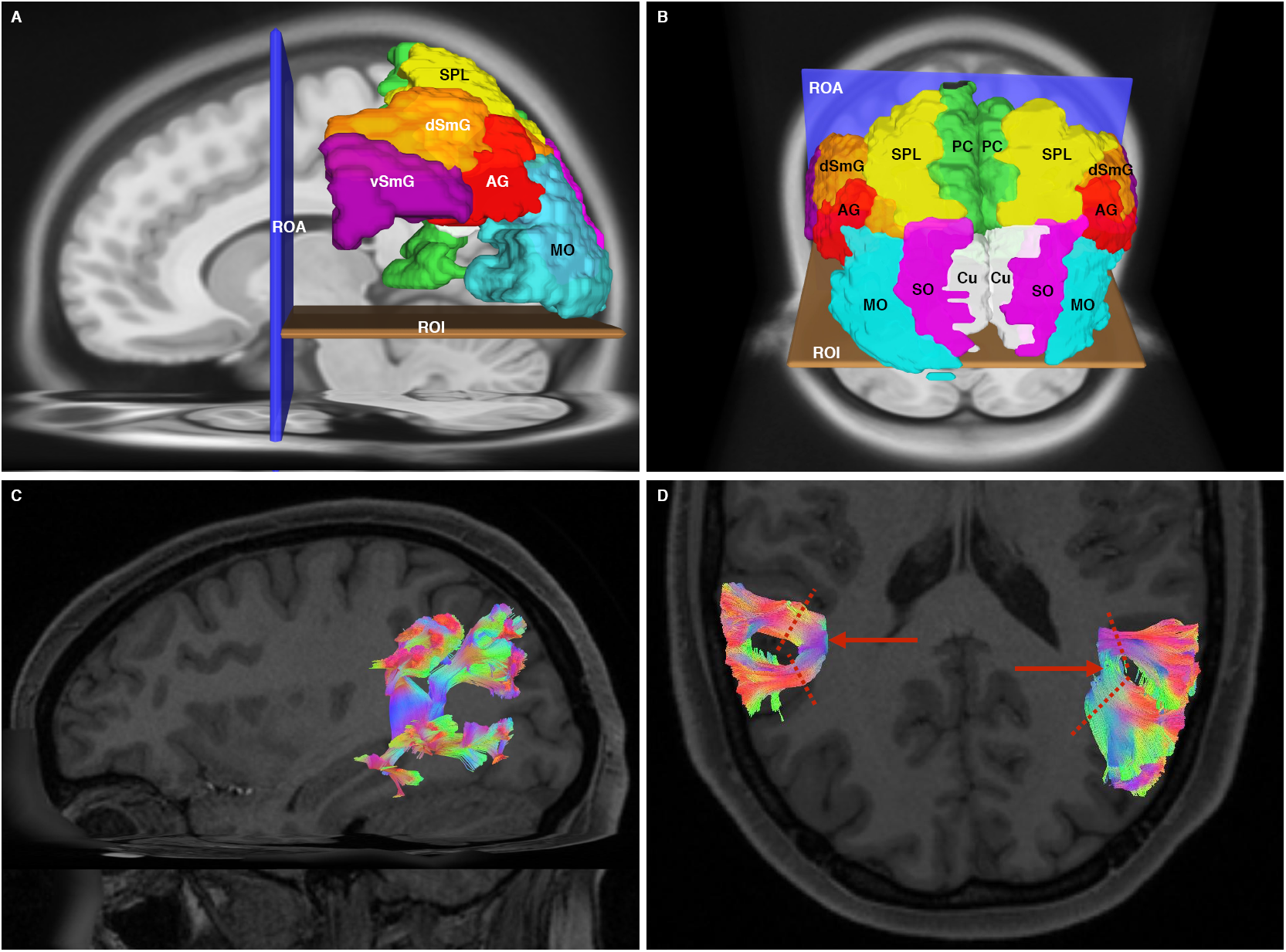
**1A** Sagittal view of the left hemisphere using the AAL atlas in the HCP 842 atlas demonstrating the seeding method for both the VAF and VOF. The ROA and ROI were the same for both bundles. For VAF supramarginal gyrus, dorsal and ventral portions (dSmG; vSmG), angular gyrus (AG), superior parietal lobule (SPL) and precuneus (PC). For the VOF, the superior occipital (SO; not visible), middle occipital (MO) and cuneus (Cu; not visible) were used. **1B** Oblique, supero-posterior view of the seeding method for the VAF and VOF within the HCP 842 atlas. The ROA and ROI were the same for both bundles. For VAF dorsal (dSmG) and ventral (vSmG) supramarginal gyrus, anguar gyrus (AG), superior, superior parietal lobule (SPL) and precuneus (PC). For the VOF, the superior occipital (SO; not visible), middle occipital (MO) and cuneus (Cu; not visible) were used. **1C** Sagittal view of the left hemisphere of a single subject, demonstrating generated VAF bundles and prior to separation. Colors are directionally assigned. **1D** Superior axial view of posterior cerebral hemispheres, demonstrating bilateral VAFs. The red arrows indicate bifurcation of the left and right bundles, respectively. The dashed red lines represent the trajectory at which the anterior and posterior bundles were subsequently selected.

**2A-D.**
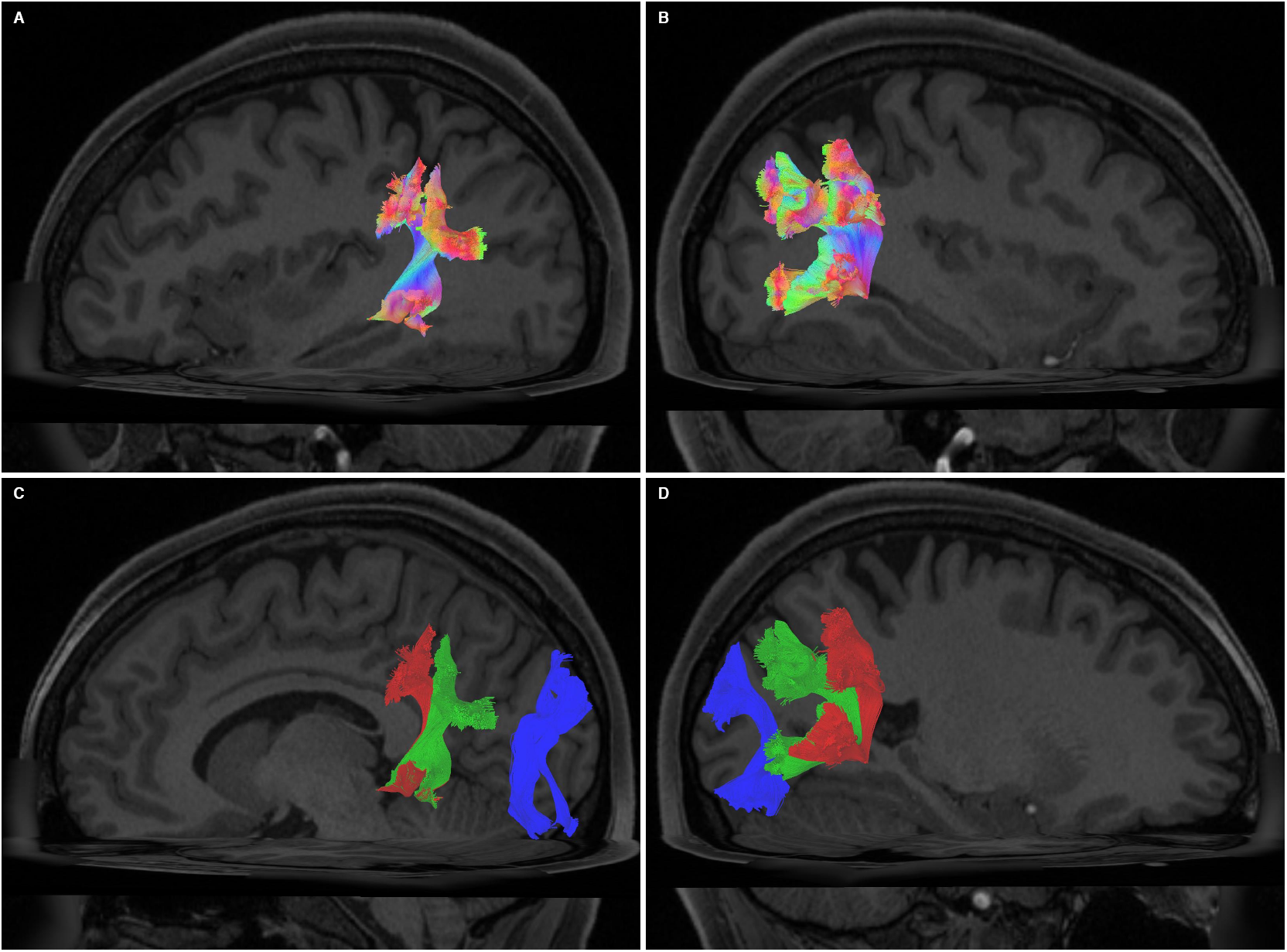
**2A** Sagittal view of the left hemisphere of a single subject (different from 1C, D) demonstrating generated VAF bundles prior to separation. Colors are directionally assigned. **2B** Sagittal view of the right hemisphere of a single subject (same as 2A) demonstrating generated VAF bundles prior to separation. Colors are directionally assigned. **2C** Sagittal view of the left hemisphere of a single subject (same as 2A). Visible are the VAF and VOF. The VAF has been segregated manually, and separate colors have been assigned to to the vVAF (red) and dVAF (green). The VOF has been colored blue. Apparent is the anteroposterior spatial separation between the entire VAF and the VOF. **2D** Sagittal view of the right hemisphere of a single subject (same as 2A). Visible are the VAF and VOF. The VAF has been segregated manually, and separate colors have been assigned to the vVAF (red) and dVAF (green). The VOF has been colored blue. Apparent is the anteroposterior spatial separation between the entire VAF and the VOF.

### 2.5 Defining Cortical Terminations

We utilized the “endpoints-to-ROI” function in DSI studio to visualize the dorsal and ventral connectivity profiles of each bundle. Once the endpoints were generated as ROI’s, these were manually compared to specific atlas regions representing various cortical gyri/Brodmann areas for connectivity analysis.

### 2.6 Quantitative Volumetry and Lateralization

We calculated the number of voxels occupied by each fiber trajectory (streamlines) and the subsequent volume (in milliliters) of each vertical-AF and VOF. Lateralization index (LI) (Catani et al., 2007) was determined using the formula ((Left tract tolume – Right tract volume) ÷ (Left tract volume + Right tract volume)) x 2. This gives an index value between −2 to +2. LI values around 0 represent general volumetric symmetry between the tracts of the left and right hemisphere. Values less than −0.4 or greater than +0.4 represent significant right or left hemispheric volumetric asymmetry, respectively. In addition to calculating LI, the volumes of left and right vertical-AFs and VOFs from the 30-subjects were each subjected to an independent samples *T*-test in SPSS (IBM Corporation, Armonk, New York) to calculate significance of mean hemispheric volumetry over the 30 subjects.

### 2.7 White Matter Fiber Dissection

Two human brain specimens were prepared for dissection. First, they were fixed in a 10% formalin solution for two months. After fixation, the arachnoid and superficial vessels were removed. The brains were subsequently frozen at −16°C for two weeks, as per the Klingler method (Ludwig & Klingler, 1956). The dissection commenced 24 hours after the specimens were thawed and proceeded in a step-wise, superficial-to-deep process. Dissection was achieved using wooden spatulas to remove successive layers of grey, and then white matter.

## 3. Results

### 3.1 Vertical Arcuate Fasciculus

#### 3.1.1 Morphology

Tracts resembling the VAF were reproduced successfully in all 30 subjects bilaterally. The fibers of the VAF assumed a ‘reverse-C’ shape form that fanned out at their parietal and temporal extremities, terminating predominantly in the middle and inferior temporal gyri and the inferior parietal gyri. From their parietal terminations, they joined approximately at the level of the Sylvian fissure into a vertically traveling stem-portion before fanning out at the temporal level (Figure 1C, D). Upon tract generation, we noticed a distinct anterior-posterior bifurcation pattern, as they traveled vertically and dorsal to the Sylvian fissure to their parietal terminations. We segmented the whole tract into two separate tracts by selecting only the fibers superior to the bifurcation. As such, we isolated two distinct tracts which henceforth are referred to as ‘ventral VAF (vVAF)’ and ‘dorsal VAF (dVAF).’ These distinct subdivisions of the VAF were each found in 58 out of 60 hemispheres (2A-D). This general morphology, including the subdivisions were found within the HCP 842 template also.

#### 3.1.2 Connectivity

We describe ventral and dorsal connectivity patterns in terms of the vVAF and dVAF. The vVAF had an anterior parietal connectivity pattern. It connected predominantly to dorsal and ventral aspects of the SmG. Connectivity of the vVAF to SmG was generally consistent in both left (83% of subjects) and right (73% of subjects) hemispheres. Dorsal SmG connectivity of the vVAF was leftward-dominant, with this connection existing in 73% of subjects left hemispheres’ but only 20% of right hemispheres. Additionally, minor vVAF connectivity to the AG was found in 30% of left hemispheres and 33% of right hemispheres.

Ventrally, the vVAF terminated within the middle temporal gyrus (MTG) in 97% and 83% of subjects’ left and right hemispheres, respectively. vVAF connectivity to the inferior temporal gyrus (ITG) was 43% and 30% on the left and right, respectively. Dorsally, the dVAF connected predominantly with the AG in 70% of subjects on the left and 93% of on the right. There was minor connectivity to ITG on both the left (20%) and right (17%). Temporal connectivity of the dVAF was hemispherically differentiated: On the left, it predominantly connected with the MTG (80%) and ITG (50%). On the right, however, it connected equally (67%) to the MTG and ITG. No connectivity of either division was found to the superior temporal gyrus (STG), or the basal occipito-temporal gyri (ITG, fusiform (FG), or lingual gyrus (LG)), (Figure 3A-B). patterns were identical within the HCP 842.

**3A-D.**
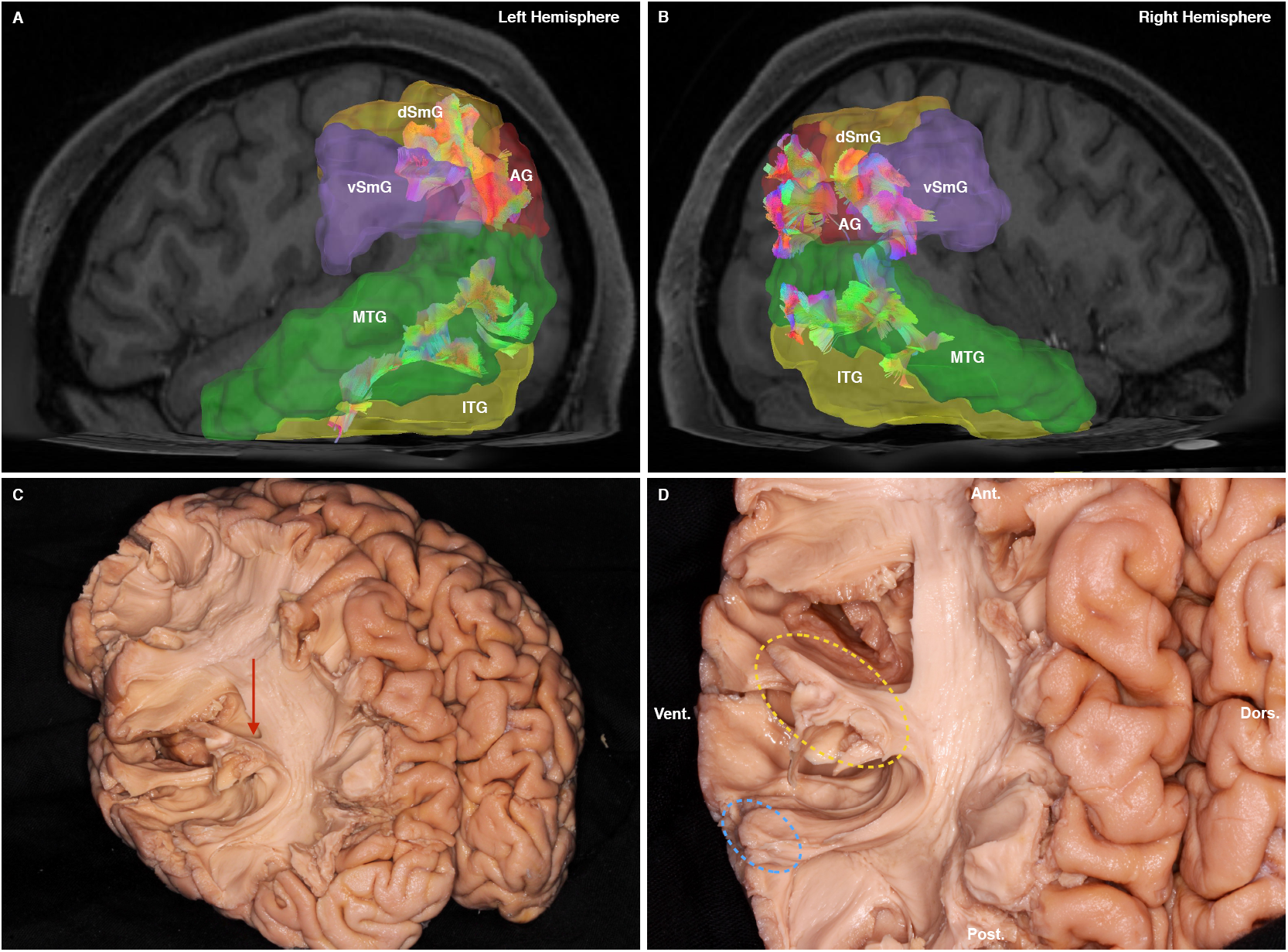
**3A** Sagittal view of the left hemisphere of a single subject. The AAL atlas regions corresponding to the dorsal (dSmG) and ventral (vSmG) portions of the supramarginal gyrus, angular gyrus (AG), middle temporal lobe (MTL) and inferior temporal lobe (ITL) are all labelled. Also apparent within the translucent, separate atlas regions are the dorsal and ventral connections of the left VAF throughout the parietal and temporal gyri. **3B** Sagittal view of the right hemisphere of a single subject. The AAL atlas regions corresponding to the relevant connections are highlighted, and are the same as in 3A. In comparison with 3A, however is the more ventral, and segregated nature of the parietal VAF connections. There are dense connections visible at the AG, and posterior ventral SmG (vSmG), with negligible connectivity to the dorsal SmG (dSmG). **3C** White fiber dissection picture of a subjects’ left hemisphere, following removal of gray matter, U-fibers and exposure of the VAF. The red arrow indicates the angular demarcation between the AF proper and the VAF, as it passes to the dorsal aspect of the sylvian fissure (i.e. inferior parietal areas). **3D** A magnified view of 3C. Orientation markers are provided at edges of the picture. The dorsal extremity of the VAF are marked by a yellow dashed-line ellipse. The ventral extremity is marked by a blue dashed-line ellipse.

#### 3.1.3 Volumetry and Lateralization

The mean volume of the left and right VAFs prior to segmentation was 9.7 ml and 9.3 ml respectively (n = 30, not significant (*t* = 0.421, *p* = 0.675)). Prior to segmenting, we used the ‘delete repeated tracts’ function in DSI studio to prevent extra tracts contributing artefactually to streamline number. After segmentation the vVAF had mean left and right hemispheric volumetry of 6.3 ml and 5.3 ml, respectively. Left and right dVAFs were smaller overall, with the right being larger than the left at 4.4 ml vs 3.8 ml. When vVAF and dVAF volumes were added, the mean volumes were 10.1 ml (0.4 ml larger than pre-segmentation) for the left VAF and 9.8 (0.4 ml larger than pre-separation) for the right VAF. Again, the difference between lateralization of the two summated VAFs was not significant (*t* = 0.470, *p* = 0.640). For non-summated VAF volumes, the LI was 0.04, and for summated volumes was 0.03, representing hemispheric symmetry. On the HCP 842 template, prior to separation the left and right VAF volumes were 5.9 and 6.9 ml on the right and left, respectively. the left vVAF was 3.9 ml and the dVAF was 2.8 ml, giving a summated volume of 6.7 ml (0.8 ml difference). On the right, volumes were 2.1 and 4.8 ml and 6.9 ml (0.1 ml difference) for the vVAF and dVAF and their sum, respectively. Lateralization indices were 0 for unmerged and −0.1 for merged VAF bundles.

### 3.2 Vertical Occipital Fasciculus

#### 3.2.1 Morphology

Vertically oriented extreme-posterior bundles resembling prior descriptions of the VOF were found in 59 out of 60 hemispheres, being absent in a single right hemisphere only. According to our results, the VOF is a ‘sheet-like’ bundle of fibers travelling from the postero-basal temporal regions and basal occipital regions to the lateral gyri of the occipital cortex. We did not find strong dorsal VOF terminations within the dorsomedial portion of the occipital lobe. Dependant on how anterior the ventral terminations of the VOF were, it assumed either an oblique dorsal and posterior trajectory, or in the case of more posterior ventral terminations, a vertical trajectory to the occipital cortex (Figure 4A, B). Using our approach, we were unable to find this short fascicle within either hemisphere of the HCP 842.

**4A-D.**
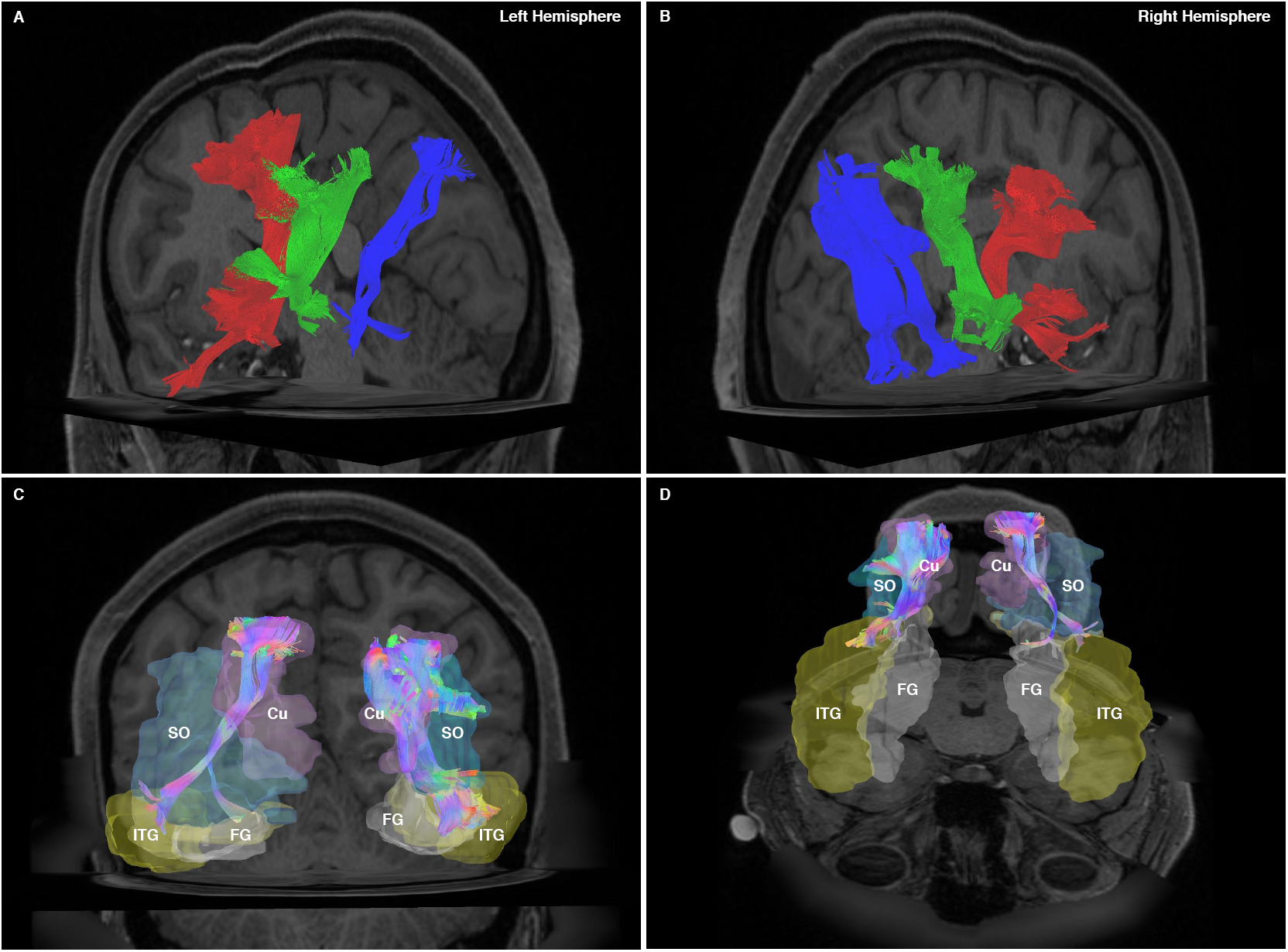
**4A** An oblique-posterior cutaway view of the left hemisphere of a single subject. Apparent are the vVAF (red), dVAF (green) and VOF (blue). From this view, the spatial separation of the VAF and VOF are readily apparent. The VOF exists as a thin sheet of fibers traversing from the basal temporo-occipital areas, to the caudal occipital areas. **4B** An oblique-posterior cutaway view of the right hemisphere of a single subject. Apparent are the vVAF (red), dVAF (green) and VOF (blue). From this view, the spatial separation of the VAF and VOF are readily apparent. The VOF exists as a thin sheet of fibers traversing from the basal temporo-occipital areas, to the caudal occipital areas. **4C** A postero-anterior view of bihemispheric, directionally colored VOFs with superimposed AAL atlas regions, from a single subject. Apparent in this diagram is the oblique, lateral-medial and ventral-dorsal trajectory of the VOF. Cu – cuneus, SO – superior portion of occipital lobe, ITL – inferior temporal lobe, Fus – fusiform gyrus. **4D** An oblique-superior, antero-posterior view of bihemispheric, directionally colored VOFs with superimposed AAL regions, from a single subject. Apparent from this view, is that the left VOF terminates within the fusiform, whereas the right VOF does not.

#### 3.1.2 Connectivity

Predominant VOF ventral connectivity was to the SO in 87% of subjects on the left, and 80% on the right. Connections to the MO were found in 37% each of subjects’ right and left hemispheres, respectively. Ventrally, VOF connectivity patterns differed hemispherically. On the left, predominant ventral connectivity was to FG (80%) and inferior occipital gyrus (IO) (60%), followed by minor connectivity to the LG (30%) and ITG (23%). On the right, connectivity was spread out relative to the left, with connectivity to FG (43%), IO (40%), ITG (30%) and LG (13%) (Figure 4C, D).

#### 3.1.3 Volumetry and Lateralization

Mean left sided VOF volume was 3.9 ml, and left-sided was 3.4 ml. Again, this difference was not significant (*t* = 0.982, *p* = 0.330). Mean lateralization index was 0.1.

#### 3.2 White Matter Fiber Dissection Results

The dissection started from the lateral surface of the hemisphere. Gyral and sulcal grey matter was removed from the precentral, postcentral, and posterior superior temporal gyri to expose underlying white matter. We preserved the gray matter of the anterior superior temporal, the majority of the middle temporal and entire inferior temporal and fusiform gyri to ensure visualization of VAF connectivity. We then exposed the dorsal aspect of the VAF at the supramarginal gyrus noting the area where the VAF and AF proper diverged approximately at the posterior extremity of the sylvian fissure. On the left, we found the VAF to originate dorsally, from the supramarginal gyrus in both brain specimens. On the right, we noted that the VAF originated slightly more posteriorly, from the region between the supramarginal and angular gyri in each specimen. Once the dorsal aspect of the VAF was exposed, we continued our dissection ventrally, removing gray and then white matter from the area below the Sylvian fissure. No connectivity of the VAF was found to the superior temporal gyrus in any hemisphere, but instead ventral terminations were at the middle temporal gyrus bilaterally, in each of the two specimens (Figure 3C, D). Once the VAF had been identified within the hemispheres, we continued our dissection posteriorly. White matter within the plane of the posterior aspect of the AF was carefully removed, to first expose the posterior oblique aspect of the middle longitudinal fasciculus (MdLF). Fibers from the caudal aspect of the parietal portion of the MdLF were carefully removed, to expose the plane containing from rostral-caudal: the posterior fan of the IFOF, the oblique VOF, and the ILF. This arrangement was observed in all 4 hemispheres (Figure 5A-D).

**5A-D.**
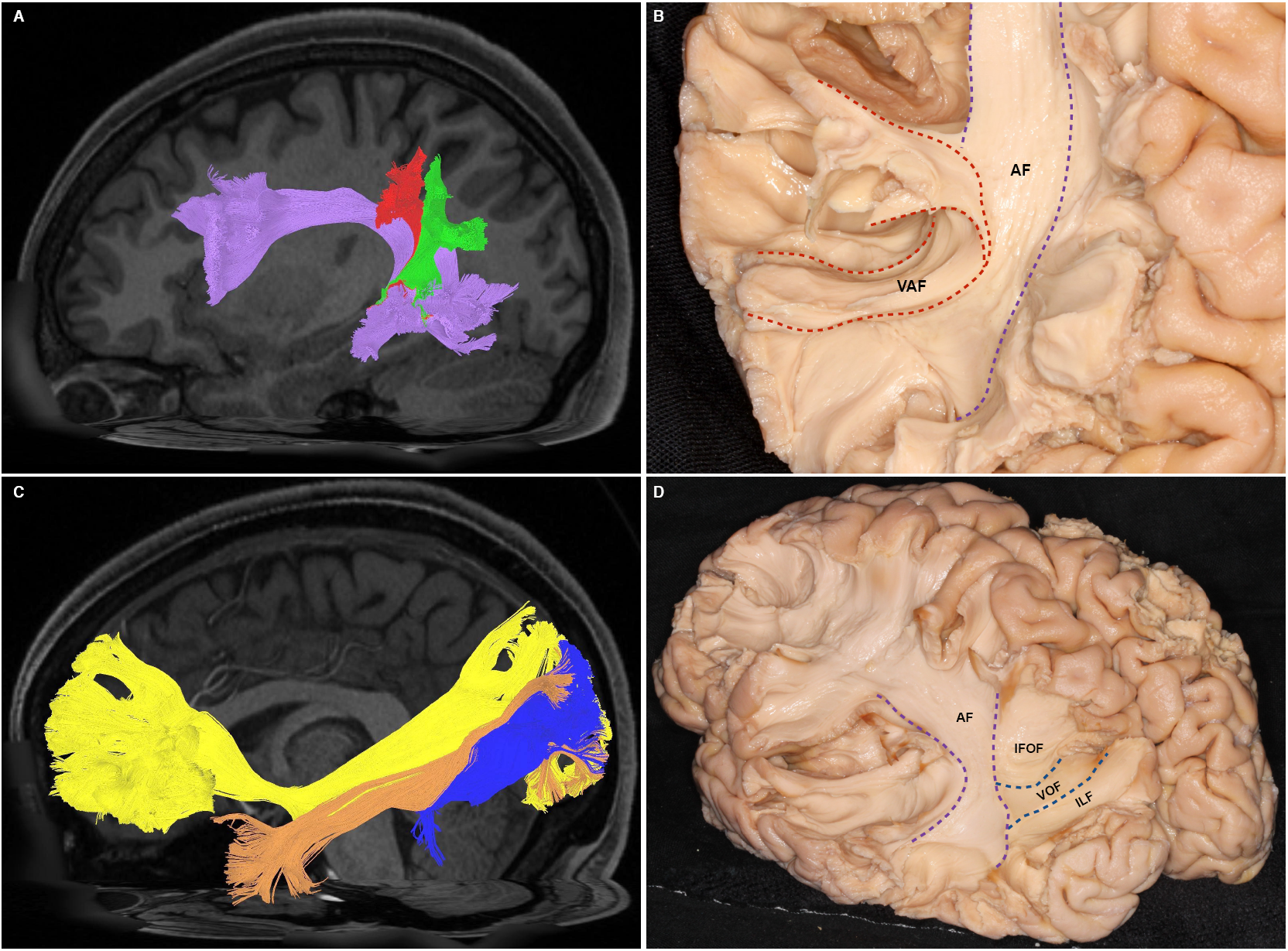
**5A** A sagittal view of the left hemisphere of a single subject. This image demonstrates the relationship of the vVAF (red) and dVAF (green) to the underlying AF (purple). **5B** A magnified view of a dissected VAF, with its demarcation from the AF proper marked out. The VAF is bounded by red-dashed lines, and the AF-proper is bounded by purple-dashed lines. **5C** A sagittal view of the left hemisphere of a single subject. This image demonstrates the superficial relationship of the VOF (blue) to the IFOF (yellow, dorsal) and ILF (orange, medial). **5D** A sagittal view of a dissected hemisphere, demonstrating the relations of the AF, posterior fan of the IFOF, VOF and ILF.

## 4. Discussion

### 4.1 Vertical Arcuate Fasciculus

Our subject-specific and template-based connectivity analysis has demonstrated consistent reproducibility of a subdivided VAF over a range of 30 single subjects, and within an average diffusion template of 842 subjects. Interestingly and uniquely, we report that this fascicle demonstrates a characteristic anteroposterior segmentation pattern. There was no significant difference between the right and left vVAF connectivity profiles, and we can infer that both are substantial conduits between the SmG and MTG, with the left vVAF also preferentially serving SMG-MTG connectivity. Likewise, the dVAF is a conduit between the AG and MTG/ITG. Broadly, our description of VAF connectivity fits in with previous descriptions of the ‘posterior-AF’ (Fernandez-Miranda et al., 2015; Wang et al., 2016).’ Our volumetric analysis revealed no indication of VAF hemispheric lateralization, instead showing that it was symmetrical. Importantly, our results can be considered further evidence that the VAF is indeed dissociated from the AF according to the ‘dorsal-ventral’ model, which emphasizes leftward-dominant lateralization of both AF components. Nor does the VAF demonstrate the rightward-dominant volumetric lateralization of the SLF system, adding further evidence to the argument for dissociation of all 3 systems due to each having distinct volumetric laterality or symmetricity.

Our connective and volumetric results indicate the VAF is an independent, short, symmetrical and vertically oriented tract running between MTG, ITG, and the entire lateral parietal lobe. The dominant vVAF, which demonstrated preferential SmG-MTG connectivity may therefore serve as a conduit for mapping sounds or read words to their meanings, the latter function potentially being subserved by MTG (Acheson & Hagoort, 2013). Furthermore, ischemia to cortex of the dominant SmG has been directly implicated in word repetition deficits (Fridriksson et al., 2010). The authors of the study implicated the ‘posterior-AF’ in observed conduction deficits. Whether they were referring specifically to the morphological entity synonymous with the VAF is uncertain, however we can infer that the VAF, and vVAF specifically is implicated in this task, from its connectivity to the inferior parietal regions. As for the dVAF, its rightward-preferential AG-MTG/ITG connectivity, may thus subserve roles in visuospatially locating objects in a 3D space. Our reasoning is based upon functional imaging studies implicating the right AG in ‘action-awareness’ (Farrer et al., 2008), and right MTG/ITG in object-location memory (Milner, Johnsrude, & Crane, 1997; Waberski et al., 2001).

Based on our data and studies into the AF and SLF, regarding nomenclature and ultimate classification of the AF amongst other close-proximity fascicles we believe the ‘perisylvian-SLF’ view to be anatomically incorrect. Firstly, an ‘arcuate-shaped tract’ cannot be at the same time a ‘vertical-shaped’ tract. Secondly, our evidence indicates segregation of AF, SLF and ‘VAF’ systems, as explained previously. Continuing to regard the AF as a ‘tripartite’ system with a vertical component defies its actual contemporary ‘dorsal-ventral’ morphology and may further confuse readers. In accordance with current terminology and in order to reconcile some controversy, we propose the term ‘parietal aslant tract’ (PAT) to more accurately describe the previously defined ‘VAF.’

### 4.2 Vertical Occipital Fasciculus

Despite lack of a concise anatomical description of an occipital vertical white matter fascicle, clinical evidence demonstrating alexia resulting from damage to occipital white matter indicated its existence. This pre-tractographic clinical data was complemented by the tractographic description of VOF anatomy (Catani et al., 2002; Wakana, Jiang, Nagae-Poetscher, van Zijl, & Mori, 2004), showing a thin sheet of vertically ascending fibers lateral to the white matter of the sagittal stratum. It wasn’t until 2013 however, that Yeatman et al. in a series of dedicated studies, offered specific elaboration of VOF structure and function. They described a tract originating from the occipitotemporal sulcus ascending vertically, lateral to the ILF and inferior fronto-occipital fasciculus, terminating in the lateral occipital and inferior parietal lobes. We elaborate upon this description with our connectivity analysis, showing a leftward-lateralized pattern of connectivity from the FG and ITG to the lateral occipital cortices (i.e. SO and MO). A 2014 functional study by (Bouhali et al., 2014) indicated leftward-lateralization of visual wordform-area (VWA) connectivity to the dorsal parieto-occipital regions. Though our results are generally congruent with this postulation, the authors of this study demonstrated that the VWA was functionally associated with perisylvian language areas. Our results do not support their latter proposition however as neither the VAF nor the VOF show FG-perisylvian connectivity.

Weiner et al. (2016) postulated that the observance of discreet VAF and VOF was attributable to tractographic artefacts, methodological issues or classification errors, concluding that they were the same tract. Our results demonstrate that the VAF and VOF are indeed distinct, spatially separated fiber bundles. Our DSI-based method of tractography specifically addresses the issue of crossing fibers, accurately differentiating them at close proximity. The vertical tracts of the posterior hemispheres can reliably be separated from abutting, perpendicular fibers, as we have demonstrated. Our results serve to refute the postulation by Weiner et al. (2016) claiming unity of the VAF and VOF. Further dissection based evidence reinforces this concept (Martino & Garcia-Porrero, 2013). In order to assure consistency of nomenclature, instead of the ‘vertical occipital fasciculus,’ we moreover propose ‘occipital aslant tract,’ as a name for this bundle and one truly representative of its structure, course and orientation.

### 4.3 Technical Considerations and Limitations

Upon attempting to generate the VOF in the HCP 842 template the reproduced fiber bundles, though resembling the VOF in morphology and trajectory at its dorsal aspects, were continuous with anteroposterior temporal fibers. These were likely false continuities. We postulate that the reason for this aberrant finding was due to the very small relative volume of the VOF. Corresponding diffusion signals from the single subjects may have been obliterated during the averaging process. As such, this is an area that must be elaborated upon, as the subject-specific analysis demonstrated the presence of this fascicle in 97% of individuals.

## 5. Conclusions

Our tractography and dissection based study into the anatomy of the VAF has yielded a novel finding that this short, vertical temporo-parietal tract can be subdivided into anterior and posterior divisions. Furthermore, we have demonstrated connective lateralization but volumetric symmetry of this system. Our evidence serves to reinforce that the VAF is a distinct fasciculus, independent of the AF and SLF systems. We therefore propose a new name for this tract: Parietal Aslant Tract, in line with current nomenclature and as an anatomical analogue to its frontal counterpart. Secondly, we have demonstrated that the parietal aslant and VOF are distinct, spatially separated systems, each demonstrating variable connectivity and thus functional specialization.

## References

Acheson, D. J., & Hagoort, P. (2013). Stimulating the brain’s language network: syntactic ambiguity resolution after TMS to the inferior frontal gyrus and middle temporal gyrus. J Cogn Neurosci, 25(10), 1664–1677. doi:10.1162/jocn_a_00430

Bouhali, F., Thiebaut de Schotten, M., Pinel, P., Poupon, C., Mangin, J. F., Dehaene, S., & Cohen, L. (2014). Anatomical connections of the visual word form area. J Neurosci, 34(46), 15402–15414. doi:10.1523/JNEUROSCI.4918-13.2014

Catani, M., Allin, M. P., Husain, M., Pugliese, L., Mesulam, M. M., Murray, R. M., & Jones, D. K. (2007). Symmetries in human brain language pathways correlate with verbal recall. Proc Natl Acad Sei USA, 104(43), 17163–17168. doi:10.1073/pnas.0702116104

Catani, M., Howard, R. J., Pajevic, S., & Jones, D. K. (2002). Virtual in vivo interactive dissection of white matter fasciculi in the human brain. Neuroimage, 17(1), 77–94.

Catani, M., Jones, D. K., & ffytche, D. H. (2005). Perisylvian language networks of the human brain. Ann Neurol, 57(1), 8–16. doi:10.1002/ana.20319

Catani, M., Mesulam, M. M., Jakobsen, E., Malik, F., Martersteck, A., Wieneke, C.,... Rogalski, E. (2013). A novel frontal pathway underlies verbal fluency in primary progressive aphasia. Brain, 136(Pt8), 2619–2628. doi:10.1093/brain/awtl63

Catani, M., & Thiebaut de Schotten, M. (2008). A diffusion tensor imaging tractography atlas for virtual in vivo dissections. Cortex, 44(8), 1105–1132. doi:10.1016/j.cortex.2008.05.004

Farquharson, S., Tournier, J. D., Calamante, F., Fabinyi, G., Schneider-Kolsky, M., Jackson, G. D., & Connelly, A. (2013). White matter fiber tractography: why we need to move beyond DTI. J Neurosurg, 118(6), 1367–1377. doi:10.3171/2013.2.JNS121294

Farrer, C., Frey, S. H., Van Horn, J. D., Tunik, E., Turk, D., Inati, S., & Grafton, S. T. (2008). The angular gyrus computes action awareness representations. Cereb Cortex, 18(2), 254–261. doi:10.1093/cercor/bhm050

Fernandez-Miranda, J. C., Rhoton, A. L., Jr., Alvarez-Linera, J., Kakizawa, Y., Choi, C., & de Oliveira, E. P. (2008). Three-dimensional microsurgical and tractographic anatomy of the white matter of the human brain. Neurosurgery, 62(6 Suppl 3), 989–1026; discussion 1026–1028. doi:10.1227/01.neu.0000333767.05328.49

Fernandez-Miranda, J. C., Wang, Y., Pathak, S., Stefaneau, L, Verstynen, T., & Yeh, F. C. (2015). Asymmetry, connectivity, and segmentation of the arcuate fascicle in the human brain. Brain Struct Fund, 220(3), 1665–1680. doi:10.1007/s00429-014-0751-7

Fridriksson, J., Kjartansson, o., Morgan, P. S., Hjaltason, H., Magnusdottir, S., Bonilha, L., & Rorden, C. (2010). Impaired speech repetition and left parietal lobe damage. J Neurosci, 30(33), 11057–11061. doi:10.1523/JNEUROSCI.1120-10.2010

Glasser, M. F., & Rilling, J. K. (2008). DTI tractography of the human brain’s language pathways. Cereb Cortex, 18(11), 2471–2482. doi:10.1093/cercor/bhn011

Lawes, I. N., Barrick, T. R., Murugam, V., Spierings, N., Evans, D. R., Song, M., & Clark, C. A. (2008). Atlas-based segmentation of white matter tracts of the human brain using diffusion tensor tractography and comparison with classical dissection. Neuroimage, 39(1), 62–79. doi: 10.1016/j.neuroimage.2007.06.041

Ludwig, E., & Klingler, J. (1956). Atlas cerebri humani: Der innere Bau des Gehirns dargestellt auf Grund makroskopischer Präparate: The inner structure of the brain demonstrated on the basis of macroscopical preparations: La structure interne du cerveau démontrée sur les préparations macroscopiques: La arquitectura interna del cerebro demostrada mediante preparaciones macroscópicas; Little, Brown.

Martino, J., da Silva-Freitas, R., Caballero, H., Marco de Lucas, E., Garcia-Porrero, J. A., & Vazquez-Barquero, A. (2013). Fiber dissection and diffusion tensor imaging tractography study of the temporoparietal fiber intersection area. Neurosurgery, 72(1 Suppl Operative), 87–97; discussion 97–88. doi:10.1227/NEU.0b013e318274294b

Martino, J., De Witt Hamer, P. C., Berger, M. S., Lawton, M. T., Arnold, C. M., de Lucas, E. M., & Duffau, H. (2013). Analysis of the subcomponents and cortical terminations of the perisylvian superior longitudinal fasciculus: a fiber dissection and DTI tractography study. Brain Struct Fund, 218(1), 105–121. doi:10.1007/s00429-012-0386-5

Martino, J., & Garcia-Porrero, J. A. (2013). Wernicke perpendicular fasciculus and vertical portion of the superior longitudinal fasciculus: in reply. Neurosurgery, 73(2), E382–383. doi:10.1227/01.neu.0000430303.56079.Oe

Milner, B., Johnsrude, L, & Crane, J. (1997). Right medial temporal-lobe contribution to object-location memory. Philos Trans R Soc LondB Biol Sci, 352(1360), 1469–1474. doi:10.1098/rstb.1997.0133

Panesar, S. S., Yeh, F. C., Deibert, C. P., Fernandes-Cabral, D., Rowthu, V., Celtikci, P.,... Fernandez-Miranda, J. C. (2017). A diffusion spectrum imaging-based tractographic study into the anatomical subdivision and cortical connectivity of the ventral external capsule: uncinate and inferior fronto-occipital fascicles. Neuroradiology. doi:10.1007/s00234-017-1874-3

Rilling, J. K., Glasser, M. F., Preuss, T. M., Ma, X., Zhao, T., Hu, X., & Behrens, T. E. (2008). The evolution of the arcuate fasciculus revealed with comparative DTI. Nat Neurosci, 11(4), 426–428. doi:10.1038/nn2072

Thiebaut de Schotten, M., Dell’Acqua, F., Forkel, S. J., Simmons, A., Vergani, F., Murphy, D. G., & Catani, M. (2011). A lateralized brain network for visuospatial attention. Nat Neurosci, 14(10), 1245–1246. doi:10.1038/nn.2905

Tzourio-Mazoyer, N., Landeau, B., Papathanassiou, D., Crivello, F., Etard, O., Delcroix, N.,... Joliot, M. (2002). Automated anatomical labeling of activations in SPM using a macroscopic anatomical parcellation of the MNI MRI single-subject brain. Neuroimage, 15(1), 273–289. doi:10.1006/nimg.2001.0978

Waberski, T. D., Kreitschmann-Andermahr, I., Kawohl, W., Darvas, F., Ryang, Y., Rodewald, M.,... Buchner, H. (2001). Spatio-temporal source imaging reveals subcomponents of the human auditory mismatch negativity in the cingulum and right inferior temporal gyrus. Neurosci Lett, 308(2), 107–110.

Wakana, S., Jiang, H., Nagae-Poetscher, L. M., van Zijl, P. C., & Mori, S. (2004). Fiber tract-based atlas of human white matter anatomy. Radiology, 230(1), 77–87. doi:10.1148/radiol.2301021640

Wang, X., Pathak, S., Stefaneanu, L., Yeh, F. C., Li, S., & Fernandez-Miranda, J. C. (2016). Subcomponents and connectivity of the superior longitudinal fasciculus in the human brain. Brain Struct Fund, 221(4), 2075–2092. doi:10.1007/s00429-015-1028-5

Weiner, K. S., Yeatman, J. D., & Wandell, B. A. (2016). The posterior arcuate fasciculus and the vertical occipital fasciculus. Cortex. doi:10.1016/j.cortex.2016.03.012

Yeatman, J. D., Weiner, K. S., Pestilli, F., Rokem, A., Mezer, A., & Wandell, B. A. (2014). The vertical occipital fasciculus: a century of controversy resolved by in vivo measurements. Proc Natl Acad Sci US A, 111(48), E5214–5223. doi:10.1073/pnas.l418503111

Yeh, F. C., & Tseng, W. Y. (2011). NTU-90: a high angular resolution brain atlas constructed by q-space diffeomorphic reconstruction. Neuroimage, 58(1), 91–99. doi: 10.1016/j.neuroimage.2011.06.021

Yeh, F. C., Verstynen, T. D., Wang, Y., Fernandez-Miranda, J. C., & Tseng, W. Y. (2013). Deterministic diffusion fiber tracking improved by quantitative anisotropy. PLoS One, 8(11), e80713. doi:10.1371/journal.pone.0080713

Yeh, F. C., Wedeen, V. J., & Tseng, W. Y. (2010). Generalized q-sampling imaging. IEEE Trans Med Imaging, 29(9), 1626–1635. doi:10.1109/TMI.2010.2045126

